# Using SNAP-tag for facile construction of dye-based biosensors in living cells

**DOI:** 10.1101/2020.07.16.206748

**Authors:** Nicholas K. Pinkin, Bei Liu, Frederico M. Pimenta, Klaus M. Hahn

## Abstract

Fluorescent biosensors based on environment-sensitive dyes have important advantages over alternative methodologies such as FRET, including the potential for enhanced brightness, elimination of bleaching artifacts, and more possibilities for multiplexing. However, such biosensors have been difficult to use because they required proteins to be covalently labeled and reintroduced into cells. Recent development of self-labeling enzymes that covalently react with membrane-permeable dyes (e.g. SNAP-tag) provide an opportunity to easily generate dye-based biosensors within cells. Here, we generate a new biosensor for Cdc42 activation by positioning SNAP-tag between Cdc42 and a peptide that binds selectively to active Cdc42. We generate a membrane-permeable Nile Red derivative that exhibits 50-fold fluorescence enhancement upon covalent labeling of the biosensor, then optimize the biosensor so the dye undergoes a 20 nm emission shift upon Cdc42 activation, enabling ratiometric imaging with a single dye. The biosensor, named SNAPsense Cdc42, is validated by examining its response to known regulatory proteins and studying Cdc42 activation during protrusion in living cells. Variants using other dyes are also presented.

## Introduction

Fluorescent biosensors have illuminated the complex interplay between localized protein activation events and cell behavior.^1–7^ Most biosensors for live-cell imaging are designed using Förster Resonance Energy Transfer (FRET) between two fluorophores whose distance or orientation is affected by the conformation of the target protein.^1,4,8^ FRET biosensors have been widely adopted by the scientific community because they are genetically encoded and therefore can be easily introduced into living cells. Unfortunately, the sensitivity of such biosensors is greatly reduced by the inefficient transfer of energy to the acceptor fluorophore. Furthermore, because the two fluorophores bleach at different rates, bleaching corrections are required for quantitative live cell imaging, and simultaneous use of multiple biosensors is hindered because two fluorophores are required for each biosensor.

These disadvantages are overcome in biosensors that are based on a single solvent-sensitive fluorophore whose fluorescence is affected by protein conformational changes.^9–16^ Such biosensors have been successful in living cells but were not frequently used because they necessitated covalent labeling of purified proteins, followed by introduction into cells through microinjection, electroporation or other cumbersome processes with a high potential to affect cell physiology.^9,12,13,15^ Biosensors have been generated in cells using unnatural amino acid (UAA) mutagenesis^17–22^, which is particularly useful because dyes can be targeted to specific amino acids. However, UAA mutagenesis requires the use of nonendogenous protein translation machinery and can be inefficient and slow to produce the biosensor.^23,24^

In the past decade, there have been significant advances in using “self-labeling” enzyme fragments, including SNAP-tag and Halo-tag, to attach dyes to proteins in living cells. These altered enzyme fragments covalently incorporate substrates bearing dyes. Labeling is efficient and fast relative to other approaches (Halo-tag, 2.4×10^6^ M^-1^s^-1^; SNAP-tag, 2.8×10^4^ M^-1^s^-1^; BCN/N_3_ strain-promoted azide-alkyne cycloaddition, 1.7 M^-1^s^-1^)^25–27^, and multiple applications have shown the method to be robust. For use of solvent-sensitive dyes to report protein activity, a potential problem is that the dye is not attached directly to the target protein, but to a ∼20-30 kDa enzyme fragment fused to the target, making it more difficult to place the dye where it will respond to target protein conformational changes.

We sought here to apply this robust intracellular labeling method to generate a biosensor based on a solvent-sensitive fluorophore, using the Rho GTPase Cdc42 to test our design strategy. Our design tethers Cdc42 to a fragment of Wiskott-Aldrich Syndrome Protein (WASP), a protein that binds selectively to the active conformation of Cdc42.^28,29^ The SNAP tag is positioned in the linker between the two, such that the dye faces the Cdc42-CBD interaction site (**Figure 1A**) and undergoes a change in environment when the WASP fragment binds Cdc42.

**Figure 1.**
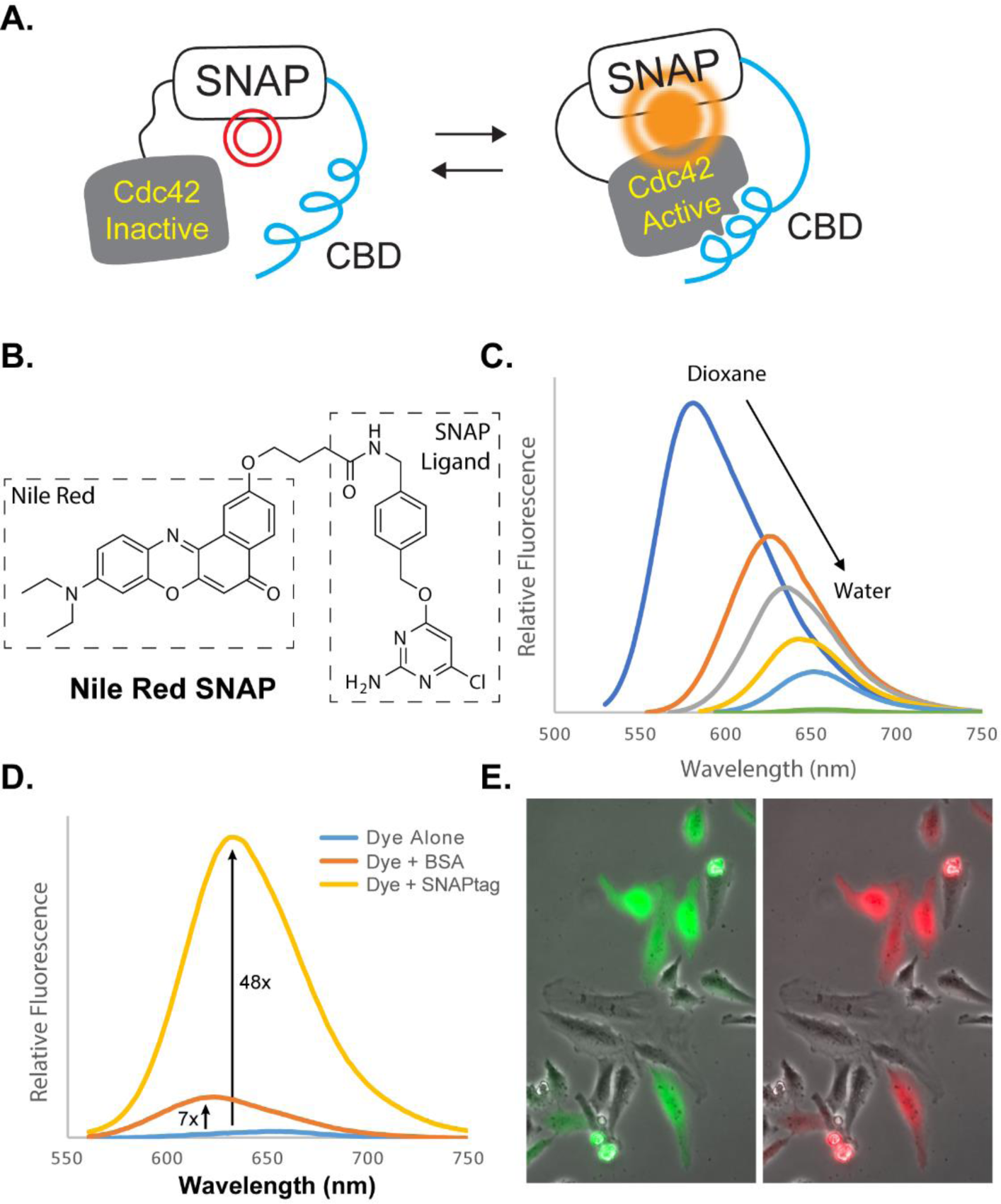
Biosensor components. (A) Design of the sensor. (B) Structure of NR-SNAP. (C) Relative fluorescence of NR-SNAP in mixtures of dioxane and water showing hypsochromic shift in emission and decrease in fluorescence as solvent polarity increases. Solvent is changed from 100% dioxane to 100% water, in 20% increments. (D) Emission increase of NR-SNAP upon attachment to SNAP-tag, compared to non-specific interaction with BSA. (E) Fluorogenic labeling. Cells expressing a GFP-tagged version of the biosensor were incubated overnight with 50 nM of NR-SNAP, followed by imaging without removing dye from the medium. NR fluorescence colocalizes with biosensor expression. Left, GFP; Right, NR.

We sought a specific set of fluorescence properties in the dye – it should increase in brightness upon covalent attachment to the probe, so that background from unattached dye would minimally affect signal/noise, and it had to be membrane permeable and not stain intracellular components. An ideal dye would undergo an environment-sensitive shift in excitation or emission maximum rather than simply an intensity change, as this would enable ratiometric imaging with a single dye. In ratiometric imaging, the readout of protein activation is the ratio of signal produced at two different excitation or emission wavelengths, eliminating effects on activation readouts from variations in cell thickness, uneven illumination or heterogeneous distribution.^30–32^ By using a single dye capable of ratiometric readout there would be no need for bleaching corrections, as each dye molecule would either contribute to the ratio or would simply be nonfluorescent. We developed a cell-permeant derivative of the solvatochromic dye Nile Red that does not stain intracellular organelles and exhibits fluorogenic turn-on labeling. After optimization, our Cdc42 sensor underwent an activity-dependent shift in emission of 20 nm, enabling ratiometric live-cell imaging using a single excitation wavelength. The biosensor responded correctly to known regulators of Cdc42 and revealed localized activation within the protrusions of motile cells.

## Results

After considering several fluorophores, we turned to the classic solvatochromic dye Nile Red (NR) because it offers a valuable combination of properties for live-cell imaging, namely large solvent-dependent spectral changes (>100 nm), long wavelength emission (∼600-650 nm), and a high extinction coefficient and quantum yield compared to other dyes with comparable solvatochromic responses.^33–36^ It has been successfully used for several applications in cells, as a lipid droplet stain^33^ and for PAINT imaging.^37,38^ Recently, proteins were derivatized with a SNAP-Tag derivative of Nile Red to follow the fluorescence changes resulting from protein-membrane interactions.^39^

We chose the Rho GTPase Cdc42 to test our SNAP-tag design strategy. Cdc42 is ubiquitous in mammalian cells, and has diverse roles including control of cell edge dynamics, apoptosis and specialized immune cell functions.^40,41^ Our previous biosensors have utilized a fragment of Wiskott-Aldrich syndrome protein, a downstream effector protein, as an affinity reagent that binds selectively to the active state of Cdc42.^28,29^ The dye is attached to the Cdc42 binding domain (CBD) of WASP at a location where binding to the target induces a fluorescence change.^9^ For our new design, Cdc42 and CBD were linked to the termini of SNAP-tag, creating a single chain construct. Adjustment of linkers positioned the CBD-Cdc42 binding interaction over the dye, so that the environment around the dye was affected by Cdc42 activation (**Figure 1A**). The termini of the SNAP tag are roughly on opposite sides of the protein, facilitating this approach. The C-terminus of Cdc42 undergoes a post-translational prenylation important for regulation,^42,43^ so we situated the Cdc42-binding domain (CBD) on the N-terminus of the SNAP-tag.

For the Nile Red SNAP-tag probe (NR-SNAP, **Figure 1B**), we utilized a short linker between NR and the SNAP ligand to ensure that the dye would not be positioned too far from the CBD-Cdc42 interaction site after labeling. NR-SNAP was non-fluorescent in water, but as the dielectric constant of the solvent decreased, it exhibited a strong hypsochromic shift and increased dramatically in brightness (**Figure 1C, Supplemental Table 1, Supplemental Figure 1**). When reacted with purified SNAP-tag protein *in vitro*, NR-SNAP underwent a nearly 50-fold increase in fluorescence (**Figure 1D and Supplemental Figure 2**) and this fluorogenic labeling was observed in living cells (**Figure 1E**).

**Figure 2.**
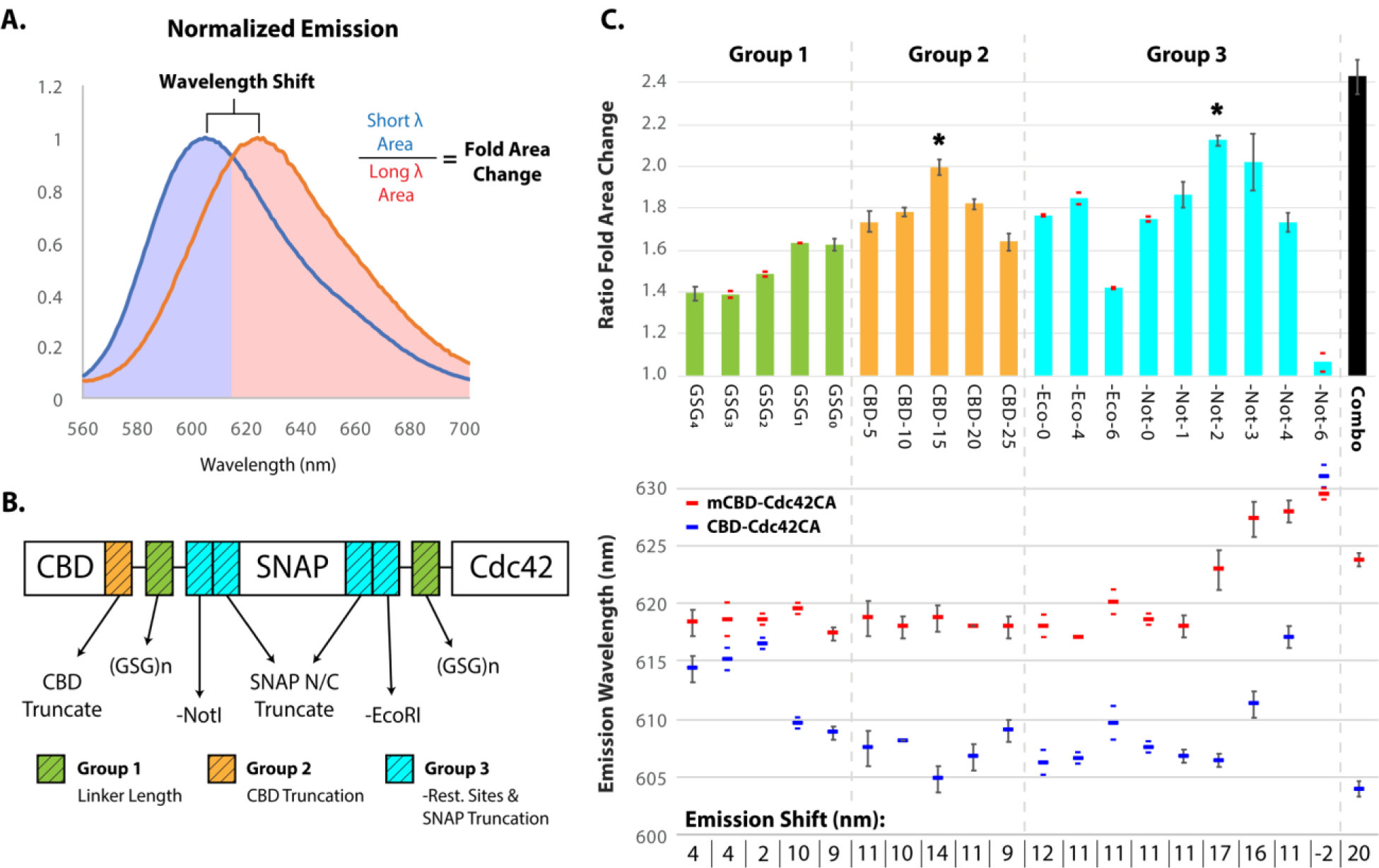
Biosensor Optimization. (A) Emission spectra of SNAPsense-Cdc42, with normalized emission intensity, for the fully optimized (combo) sensor, illustrating the emission shift and the portions of the spectra used to calculate the fold area change. Blue curve is the positive control (CBD-Cdc24CA) and red curve is the negative control (mCBD-Cdc42CA). (B) Sensor design and regions of focus for optimization. (C) Measured fold area changes and emission shifts for each variant of the sensor tested. Stars indicate the optimal modifications from each round that were combined to produce the fully optimized construct on the far right. Numbers at the bottom are the wavelength shifts (nm) between the positive and negative controls. Data are plotted as the mean ± s.d., n = 3 biologically independent samples when error bars are shown. For all other measurements, n = 2 biologically independent samples (individual ratio values and measurements are shown above and below the mean).

To optimize the extent of fluorescence change induced by Cdc42 activation, we compared biosensors with different point mutations. A Q61L mutation^44^ was used to produce constitutively active Cdc42 (CBD-Cdc42CA), used as a positive control. For a negative control, Cdc42 Q61L was combined with CBD containing mutations that greatly reduce Cdc42 affinity, H246D & H249D (mCBD-Cdc42CA).^9^ We used HeLa cells for all optimization studies because they robustly expressed the biosensor across much of the cell population. The NR emission spectrum from cells in suspension were measured for each construct. Optimization was based on two parameters: the shift in maximum emission wavelength and the ratio of the areas under two portions of the emission spectra, chosen to mimic filters used for ratiometric imaging in live cells (**Figure 2A**).

We began by testing various lengths of flexible and soluble Gly-Ser-Gly (GSG) linkers as spacers between each of the sensor components. The dynamic range of the sensor improved as the linker length decreased (**Figure 2 B & C**, Group 1), consistent with the idea that shorter linkers would orient the CBD-Cdc42 complex closer to the dye. Given the trend toward improved response with shorter linkers, we reasoned that we should try truncating the proteins themselves at the interfaces. We chose not to truncate Cdc42 at all to ensure that there would be no effect on activity and biological interactions. A large portion of the C-terminal sequence of CBD is not involved in the interaction with Cdc42,^29^ so we tested the truncation of the last 25 amino acids in 5 amino acid intervals (**Figure 2 B & C**, Group 2). This led to increasing dynamic range when removing up to 15 amino acids (CBD-15), then a decrease as more amino acids were removed, perhaps because the more extensive truncations prevented the protein fragments from reaching one another. Due to the extensive number of experiments needed for optimization, some values were tested in triplicate and some in duplicate, as indicated.

We next explored truncating the N- and C-termini of the SNAP-tag, as well as removing the EcoRI and NotI restriction sites used at each terminus for cloning (**Figure 2 B & C**, Group 3). Though short, linkers encoded by the restriction sites could diminish the response of the sensor. At the N-terminus, we observed a slight increase in the dynamic range when EcoRI was removed, and further increases when SNAP-tag was truncated by two or four amino acids. Removal of six amino acids largely abolished dye attachment, likely because this portion includes a Cys that binds Zn^2+^ involved in stabilizing the SNAP-tag^44^ (although an activation-dependent response was still observed). At the C-terminus, removing NotI and truncating one amino acid from SNAP-tag led to small improvements in dynamic range, but removal of additional amino acids had significant effects on the dynamic range, and for the first time produced a red-shift of the emission maximum for the off state. Truncating two amino acids red-shifted the inactive state by 7 nm without affecting the on state, resulting in an 18 nm overall shift and a 2.1-fold change in area ratio (**Figure 2C**, Group 3, Not-2). Further truncations continued to red shift both states of the sensor, decreasing the dynamic range of the sensor and ultimately abolishing the response when six amino acids were removed (though the labeling reaction was unaffected). Combining the best modifications at the N- and C-termini of SNAP-tag (CBD-15, -NotI/-2 AA SNAP C-terminal truncation) provided us with our optimized sensor, which exhibited a 20 nm emission shift and a nearly 2.5-fold change in area ratio. In all experiments described below we used this optimized construct, named simply the SNAPsense-Cdc42 biosensor.

It was important that the new biosensor respond appropriately to upstream signals in living cells, as it is in effect an indicator of how these signals converge on Cdc42. We sought to validate that the optimized biosensor responds to both activating guanine exchange factors (GEFs) and the negative regulator RhoGDI (**Figure 3, A & B**). For this, we generated a version of the biosensor containing a dominant negative mutation in Cdc42 (SNAPsense-Cdc42DN, T17N)^45–47^ and compared its response in cells to the wild type version (SNAPsense-Cdc42WT) and to the positive and negative controls described above. As expected, the WT biosensor exhibited a response intermediate between the positive and negative controls, indicative of a mixture of active and inactive Cdc42 within cells. The DN biosensor was slightly red shifted and had a lower area ratio compared to the WT biosensor, but did not respond as strongly as the negative control. The weak binding of the T17N dominant negative mutant of Cdc42 to CBD may be enhanced in this control biosensor because the two proteins are undergoing an intramolecular interaction.

**Figure 3.**
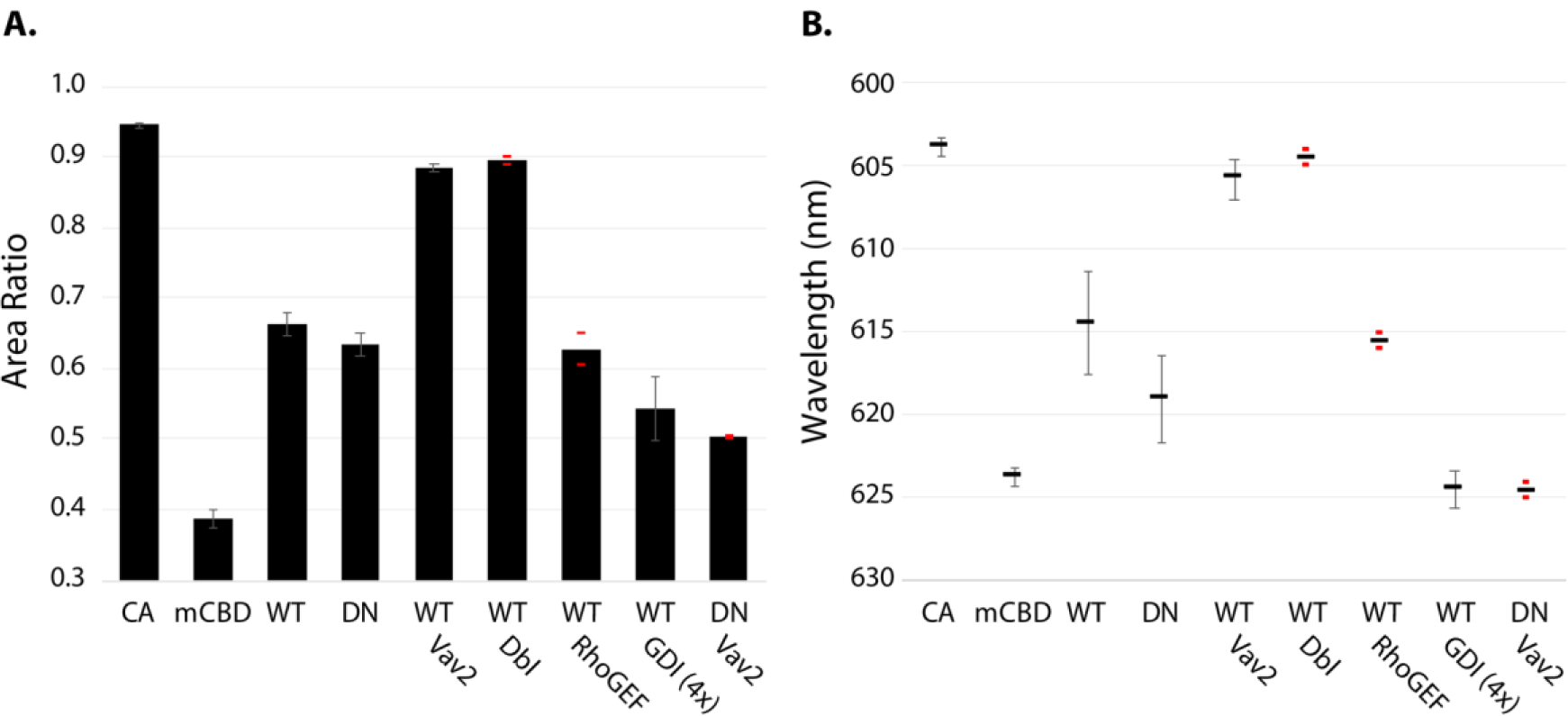
Biosensor response to regulatory molecules. (A,B) Fluorescence response of SNAPsense-Cdc42 co-expressed with activating GEFs (Vav2 and Dbl), a GEF not specific for Cdc42 (RhoGEF), and the negative regulator RhoGDI. Data are plotted as the mean ± s.d., n = 3 biologically independent samples when error bars are shown. For all other measurements, n = 2 biologically independent samples (individual measurements are shown as red marks above and below the mean).

Co-expression of SNAPsense-Cdc42WT with the Cdc42-specific GEFs Vav2 and Dbl produced a blue shift and an increase in the area ratio to generate fluorescence like that of the CA control. In contrast, co-expression of the RhoA-specific GEF p115-RhoGEF had little effect, and co-expression of RhoGDI decreased the area ratio and red-shifted the fluorescence to match the negative control (see also Supplemental Figure 3). Interestingly, co-expression of Vav2 with the DN sensor also red-shifted the emission. GEFs bind preferentially to both the nucleotide-free and GDP-bound states of GTPases, and RhoGDI functions to bind to GTPases and sequester them from the membrane.^43,48–50^ These binding interactions inhibit the binding of Cdc42 by CBD, reflected by the red-shift we observe, and this effect is pronounced for the DN sensor likely because it cannot be converted to the GTP-loaded state by Vav2. Our results demonstrate that fluorescence of SNAPsense-Cdc42WT reflects an intermediate level of Cdc42 activation in living cells, and that the biosensor responds to stimuli that shift this activation equilibrium to more active Cdc42 (GEFs) or to inactive Cdc42 (RhoGDI).

Finally, we applied SNAPsense-Cdc42 to examine the localization and kinetics of Cdc42 activity in living Mouse Embryonic Fibroblasts (MEFs) undergoing constitutive protrusion/retraction cycles. Multiple studies have shown that Cdc42 plays an important role in cell protrusion, and that interaction with other GTPases and effector proteins is spatially and temporally controlled.^8,9,44,51–55^ We generated a cell line stably expressing SNAPsense-Cdc42WT with biosensor expression under the control of a doxycycline-regulated promoter.^56^ This proved substantially easier than using our previously published Cdc42 biosensor in which dye is incorporated through UAA labeling ; this requires co-expression of the UAA-specific tRNA synthetase and multiple copies its orthogonal tRNA. The biosensor was excited with a 561 nm laser and emission was split by a dichroic passing the wavelengths above and below 624 nm to different cameras (Semrock, FF624-Di01). After background subtraction and registration, the “short” wavelength image was divided by the “long” wavelength image to produce a ratio image reflecting Cdc42 activity. Because the dye underwent an emission shift, and not simply a change in intensity, ratio imaging could be accomplished without the need to attach a second fluorophore (e.g. a fluorescent protein) to normalize dye changes for variations in illumination intensity and cell thickness.

The ratio values within cells expressing SNAPsense-Cdc42WT were substantially lower than those in cells with constitutively active biosensor (SNAPsense-Cdc42CA) (**Figure 4A & B, Supplemental Movie 1**). This was apparent in observation of ratio images and was quantified by examining histograms of ratio values across cell populations (**Figure 4C**). The constitutively active control biosensor showed high activation across the entire cell, and prevented the normal broad extensions seen in MEFS (**Figure 4B**). The WT biosensor showed increased activity in extending protrusions (**Figure 4A**). To quantify this, we used the program Edgeprops to examine the relationship between cell edge velocity and nearby Cdc42 activity levels (**Figure 4D**).^57^ This showed a roughly linear increase in ratio value with protrusion velocity, and lower activation levels during retraction.

**Figure 4.**
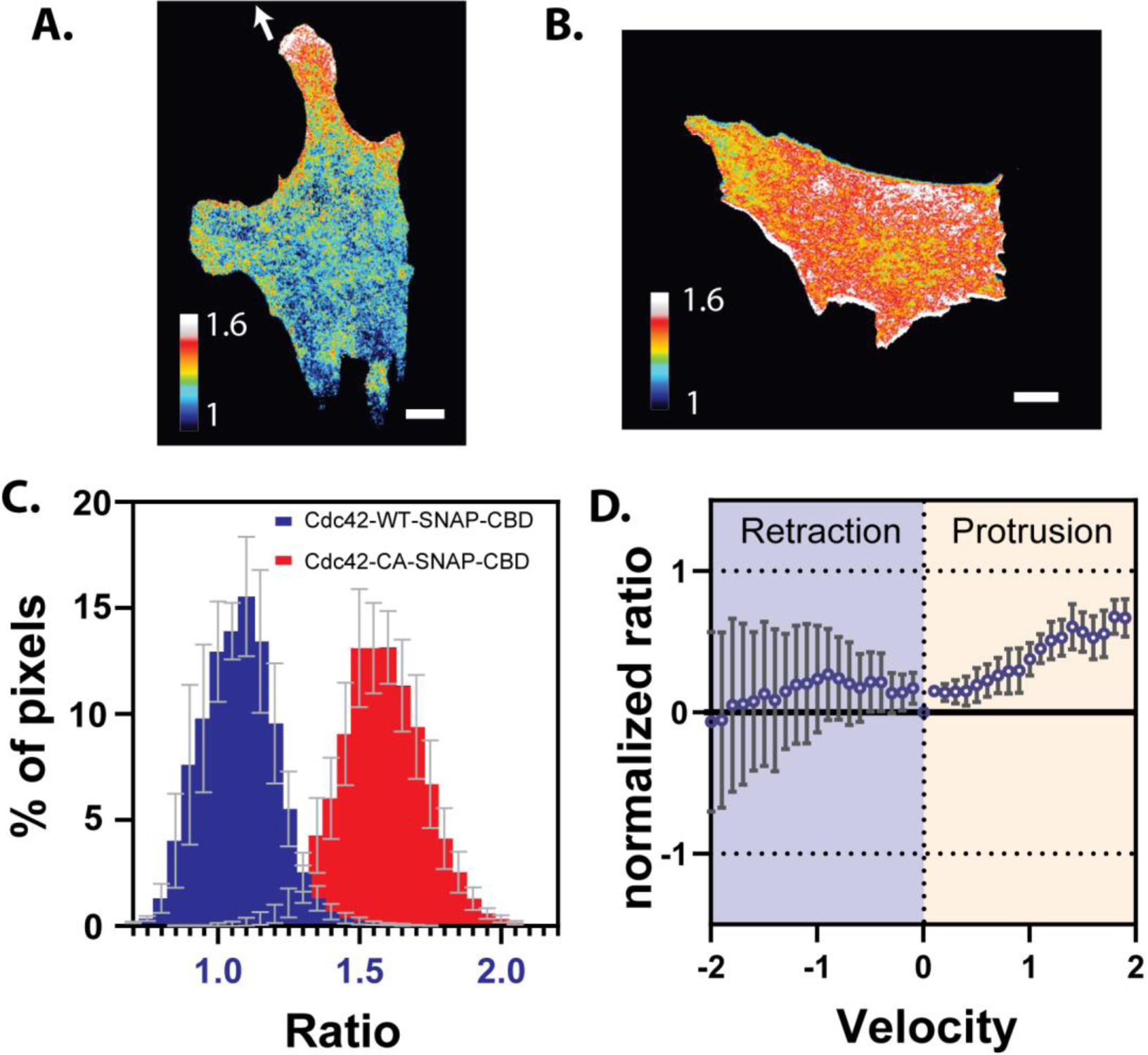
Cdc42 activity in living cells. (A.) TIRF imaging of MEF cells expressing CBD-SNAP-Cdc42WT, labeled with NR-SNAP. Cdc42 activity is seen in cell protrusions. (B.) MEF cells expressing CBD-SNAP-Cdc42CA showed high activity across the whole cell, with higher levels at the cell edge. (C.) Histogram of the pixel-by-pixel ratio distribution of Cdc42-WT (6 cells) and Cdc42-CA (10 cells). Data are plotted as the mean ± s.e.m. (D.) Plot of Cdc42 activation versus cell edge velocity analyzed by EdgeProps (6 cells).^57^ Data are plotted as the mean ± s.e.m. (scale bar = 10 μm, color scales indicate relative ratio values, excluding upper and lower 5% of pixels. Images in A and B are scaled identically)

Our SNAPsense biosensor is a simple design that capitalizes on the hypsochromic shift of Nile Red to produce a ratiometric fluorescence change using only one fluorophore. However, to enhance biosensor sensitivity it could be valuable to take advantage of bright dyes that undergo large environment-dependent changes in fluorescence intensity.^15^ We therefore appended superfolder GFP (sfGFP) on the N-terminus of the biosensor (**Figure 5A**) to act as an unchanging fluorophore for normalization of an intensity-changing dye. Appending sfGFP had no effect on the spectral response of Nile Red, indicating that the N-terminus can be flexibly modified (**Figure 5B**). NR normalization against sfGFP allowed us to gauge intensity changes in cell suspensions despite heterogenous expression levels. The NR biosensor reflected activation with a 2-fold change in brightness, in addition to the 20 nm emission shift. The brightness change and spectral shift worked in tandem to increase dynamic range. Interestingly, incorporation of the SNAP ligand onto a different portion of Nile Red decreased the brightness change and increased the spectral shift (**Figure 5B**), suggesting that the donor and acceptor portions of the dye interact differently with the protein. Such information could be helpful for the further rational improvement of the biosensor.

**Figure 5.**
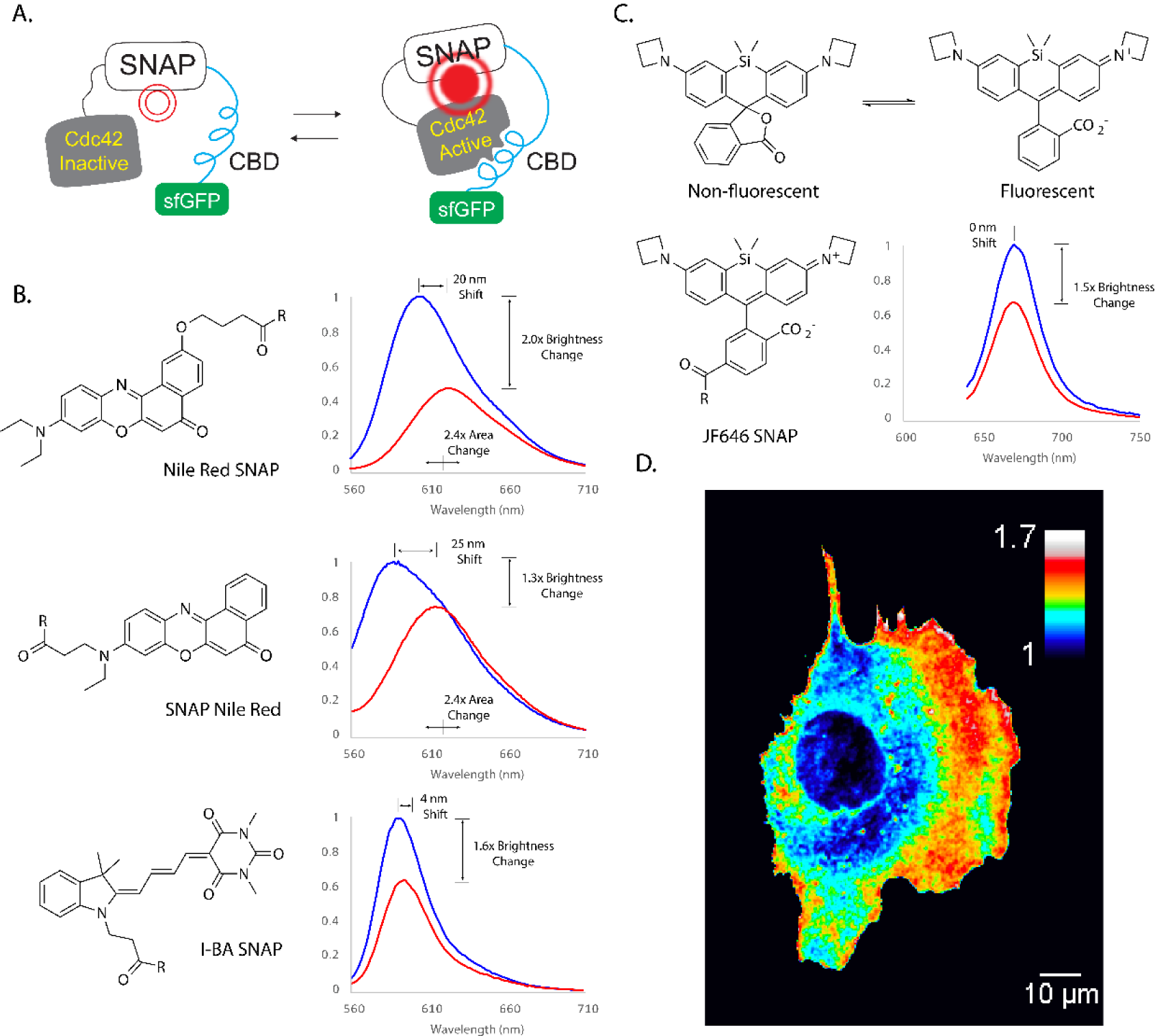
Biosensors with alternate environment-sensitive dyes. Addition of sfGFP to Cdc42 SNAPsensor for ratio imaging with dyes that undergo intensity changes. (A) The N-terminus of CBD modified with the fluorescent protein sfGFP. (B) top and middle: moving the reactive side chain to incorporate Nile Red with different orientations; bottom: The merocyanine dye I-BA showed an activation-dependent change in fluorescence intensity. (C) Rhodamine dyes exist in an equilibrium between a non-fluorescent lactone and a fluorescent zwitterion, which is the basis for the fluorogenic response when labeling SNAP-tag or Halo-tag. Cdc42 led to increased activation, possibly because the change in dye environment affected the equilibrium. Fluorescence data are representative data from n = 2 biologically independent samples. (D) Imaging Cdc42 activity in a MEF cell using dye JF646 (Scale as in Figure 4). *I-BA and JF646 were tested on multiple versions of the biosensor generated during optimization for NR-SNAP; data is shown for the biosensor that produced the maximum response for each dye (see materials and methods).

We used the sfGFP-tagged biosensor to compare the environment-dependent changes of two other dyes incorporated into the biosensor through SNAP-tag conjugation, the merocyanine dye I-BA, and the rhodamine-derivative JF646 (**Figure 5 B**,**C**).^11,58^ As expected, I-BA responded by increasing in brightness with little change in emission wavelength. For this dye, the greatest response occurred using the CBD-15 version of the biosensor (**Figure 2C**, group 2). Rhodamine dyes exist in an equilibrium between a fluorescent zwitterionic form and a non-fluorescent, neutral lactone form.^58–60^ This leads to fluorogenic labeling of SNAP-tags and Halo-tags. Given that labeling can force the equilibrium to favor the fluorescent state, we wondered whether binding between CBD and Cdc42 in the biosensor could modulate the equilibrium and thereby the dye brightness. Indeed, JF646-SNAP showed a 1.5-fold increase in brightness on the GSG_0_ sensor (**Figure 2C**, group 1; **Figure 5C**). Interestingly neither I-BA nor JF646 showed activation-dependent changes on the biosensor optimized for NR-SNAP, but required re-optimization of the spacing between biosensor components, indicating that specific interactions between protein and dye components play an important role in the response.

Using a MEF cell line stably expressing the sfGFP-tagged GSG_0_ sensor we were able to image Cdc42 activity using the JF646 dye for ratio imaging. Activation was observed in cell protrusions as described for the Nile Red biosensor (**Figures 4 and 5D**). This proof of concept example illustrates the flexibility of our SNAP-tag design for use with different dyes. While sfGFP was used for this example, the fluorescent protein might be replaced with HaloTag to enable a far-red dye to be introduced as a reference for ratio imaging. Such a design could be useful for multiplexing our Cdc42 sensor with other FRET biosensors, or potentially for imaging in more complex samples such as tissues or living organisms, where longer wavelengths are necessary.

## Conclusion

Our SNAP-tag biosensor for Cdc42 represents a new type of biosensor design that can greatly simplify the use of fluorescent dyes to study protein activity in living cells. We believe it is an attractive alternative to FRET because it harnesses the advantages of environment-sensing dyes to provide bright, ratiometric readouts without the need for bleaching correction, and will extend the ability to make measurements over a broad range of wavelengths. Although use of a single dye is valuable for these advantages, we have also shown that a second dye can be incorporated to enable ratio imaging with dyes that undergo intensity changes. This provides access to dyes that respond to different environmental parameters, and to bright environment-sensing dyes that fluoresce over a broader range of wavelengths. The use of far red dyes will be valuable for biosensor imaging in tissues or live animals.

In the future, we hope this approach will benefit single molecule applications, multiplexing through the use of different orthogonal labeling approaches in the same cell, and exploring the biology of low abundance proteins made accessible because bright dyes can be used at lower expression level.

## Acknowledgements

We thank Luke Lavis (Janelia Research Campus) for providing Janelia Fluor dyes and Mauro Calabrese (University of North Carolina at Chapel Hill) for vectors used to generate stable cell lines. We gratefully acknowledge the National Institute of General Medical Sciences of the National Institutes of Health (N.K.P., F32GM120958 and K.M.H., R35GM122596) for financial support of this research.

