## Supplemental Information for "Using SNAP-tag for facile construction of dye-based biosensors in living cells"

### Supplemental Info

#### General materials and methods

All reagents and solvents were purchased from commercial suppliers (TCI America, Acros, Sigma Aldrich, Fisher Scientific) and used without further purification. TLC chromatography was performed using TLC Silica gel 60G F254 25 Glassplates (EMD Millipore). UV-visible spectra were obtained with a Hewlett-Packard 8453 diode array spectrophotometer. Chromatography was carried out on an Isco Combiflash Companion apparatus. Emission and excitation spectra were obtained using a Horiba Fluorolog 3 spectrofluorometer at room temperature. Mass spectra were obtained on a ThermoScientific Q Exactive HF-X using direct infusion. NMR spectra were obtained on an Inova-400 spectrometer. Chemical shifts are reported using NMR reference peaks of 7.26 ($^1$H) and 77.23 ($^{13}$C) ppm for $CDCl_3$, and 2.50 ($^1$H) and 39.51 ($^{13}$C) ppm for DMSO-$d^6$.

#### Synthesis of Nile Red SNAP

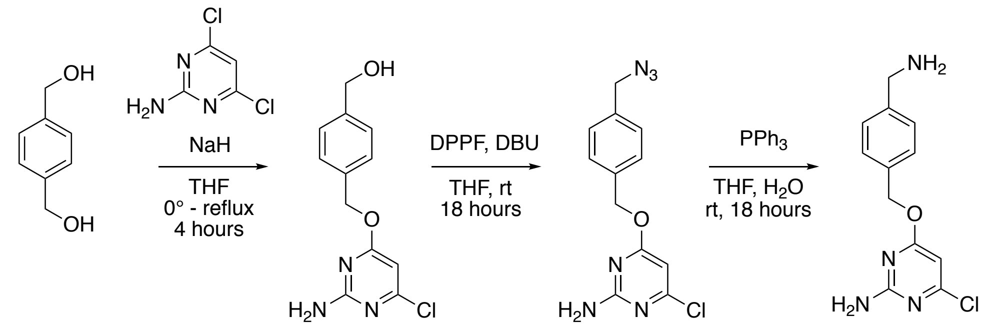


1 2 **SNAP-NH$_2$**

**Supplemental Scheme 1.** Synthesis of SNAP-$NH_2$.

**(4-(((2-amino-6-chloropyrimidin-4-yl)oxy)methyl)phenyl)methanol (1).** Benzene dimethanol (3.37 g, 24.4 mmol) and 4-Amino-2,6-dichloropyrimidine (1.0 g, 6.1 mmol) were dissolved into dry THF (11 mL) under argon and cooled to 0 °C. Sodium hydride (60% dispersion in mineral oil, 234 mg, 6.1 mmol) was added in one portion and the reaction was allowed to continue to stir at 0 °C for 10 minutes. Then the reaction was brought to reflux for four hours. The reaction was cooled to room temperature and quenched with methanol, then was dried onto silica gel. The product was purified on silica gel using a gradient of 0-10% methanol in DCM to yield a yellow solid (1.21 g, 80% yield).

$^{1}H$ NMR (400 MHz, DMSO-$d_6$) δ 7.39 (d, *J* = 8.1 Hz, 2H), 7.32 (d, *J* = 8.1 Hz, 2H), 7.10 (s, 2H), 6.14 (s, 1H), 5.30 (s, 2H), 5.18 (t, *J* = 5.7 Hz, 1H), 4.49 (d, *J* = 5.6 Hz, 2H). $^{13}C$ NMR (100 MHz, DMSO-$d_6$) δ 170.33, 162.79, 160.00, 142.53, 134.49, 128.17, 126.43, 94.42, 67.34, 62.61. HRMS (M + $H^+$): Expected: 266.0691, Found: 266.0681.

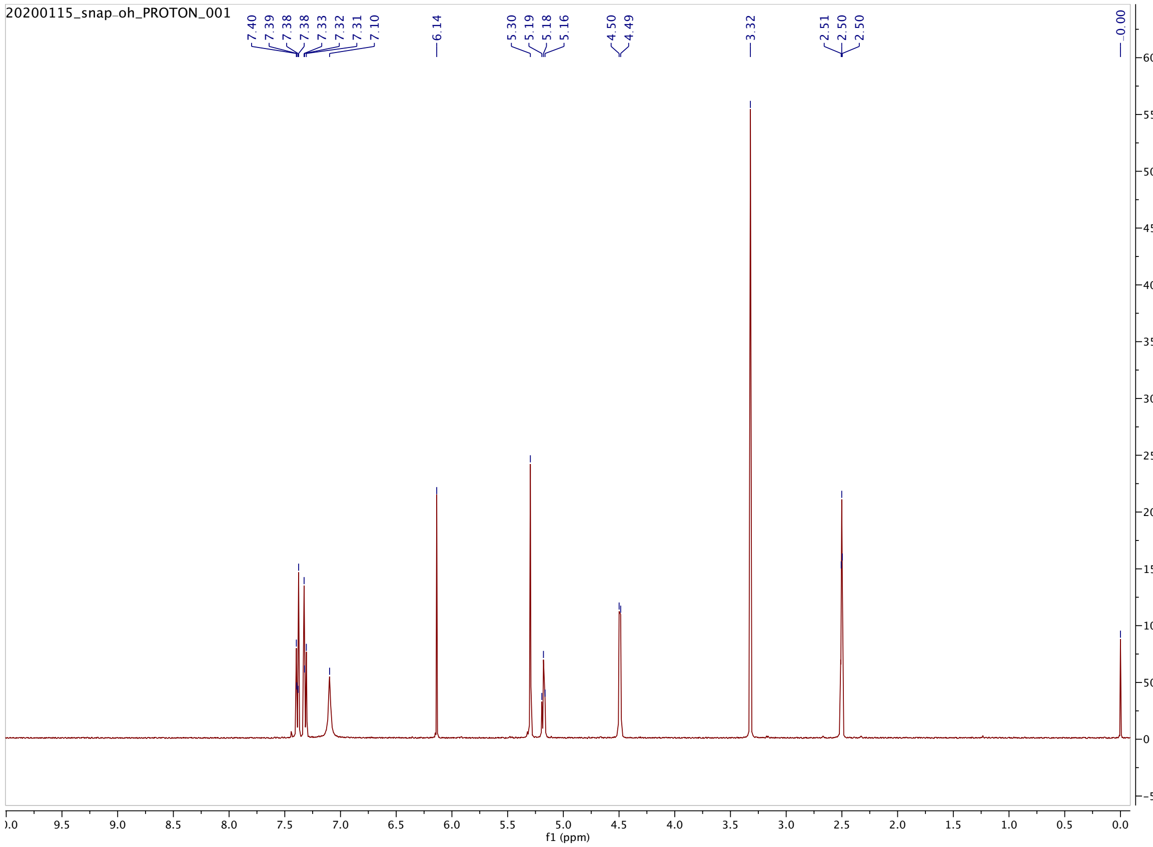

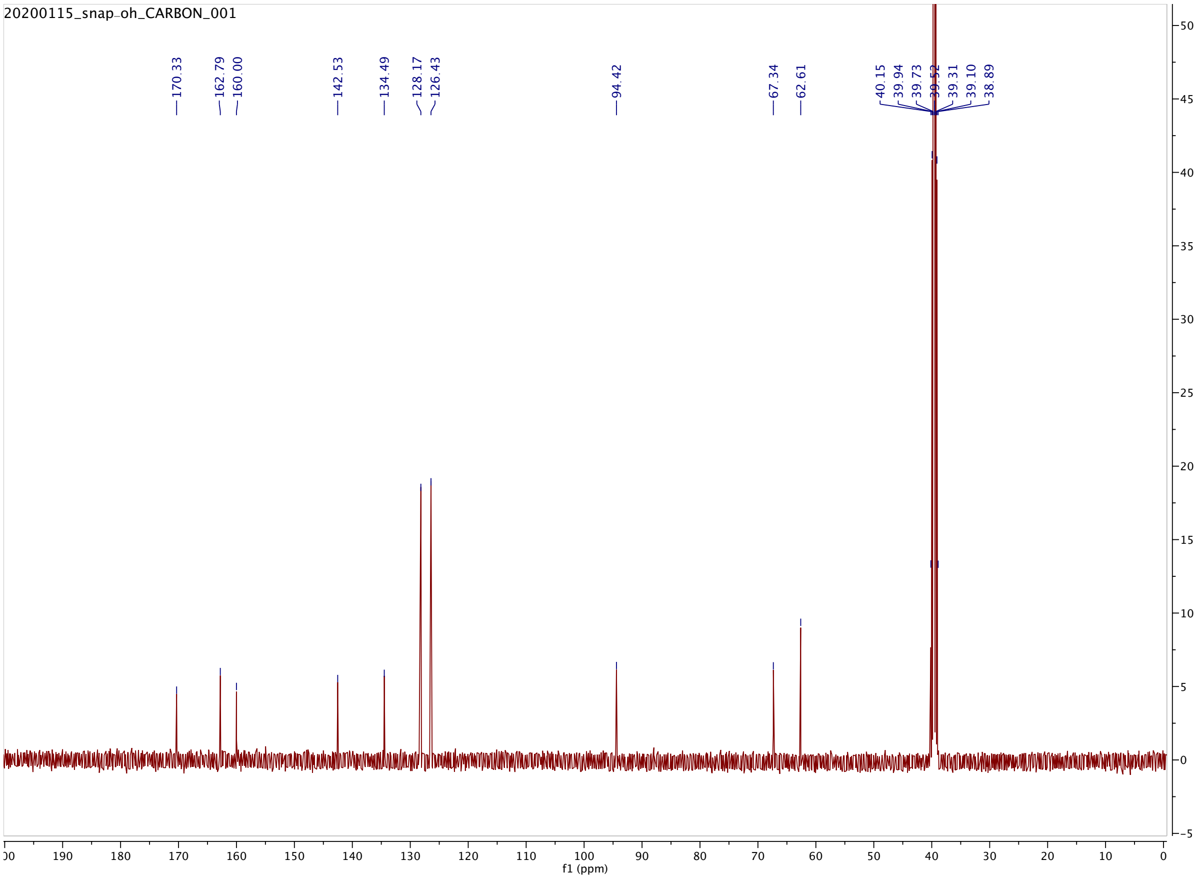

20200115_snap-oh_CARBON_001
170.33
162.79
160.00
142.53
134.49
128.17
126.43
94.42
67.34
62.61
40.15
39.94
39.73
39.52
39.31
39.10
38.89
f1 (ppm)

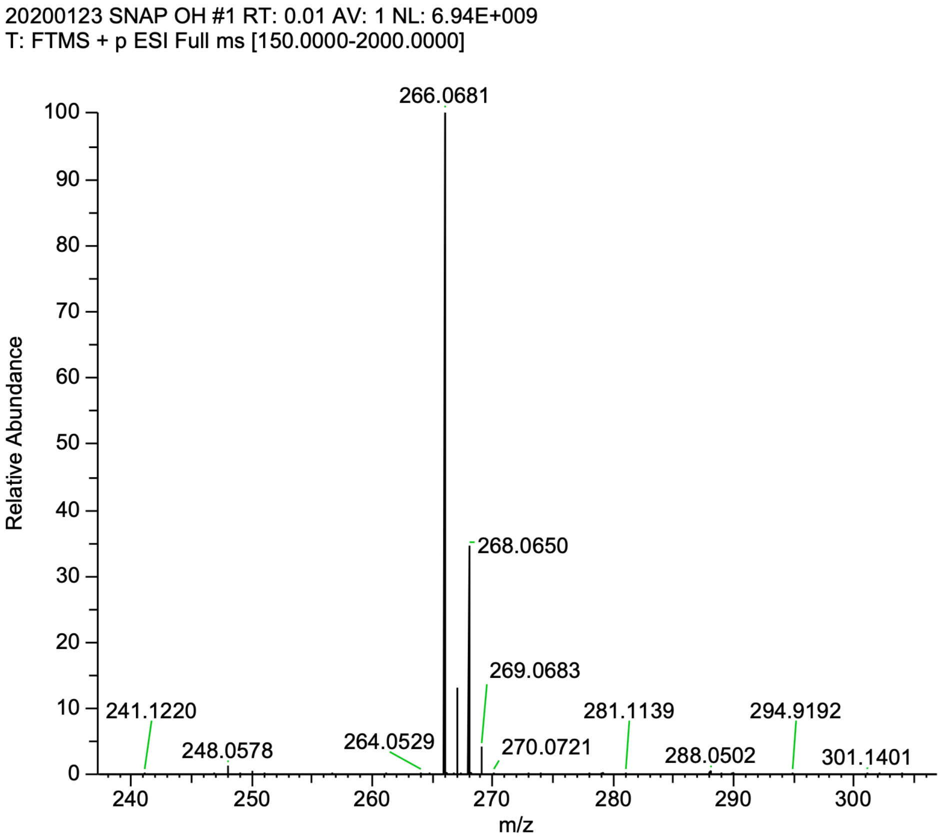


**4-((4-(azidomethyl)benzyl)oxy)-6-chloropyrimidin-2-amine (2). 1** (500 mg, 1.85 mmol) was dissolved into dry THF (38 mL) under argon, then DBU (0.414 mL, 2.78 mmol) and diphenylphosphoryl azide (DPPA, 0.476 mL, 2.21 mmol) were added. The reaction was allowed to stir at room temperature overnight, after which TLC showed complete conversion to product. The reaction solution was dried onto silica gel and the product purified with a gradient of 10-40% ethyl acetate in hexanes to yield a fluffy white solid (503 mg, 99% yield).

$^1$H NMR (400 MHz, DMSO-$d_6$) δ 7.47 (d, 2H), 7.39 (d, 2H), 7.11 (s, 2H), 6.16 (s, 1H), 5.33 (s, 2H), 4.45 (s, 2H). $^{13}$C NMR (100 MHz, DMSO-$d_6$) δ 170.28, 162.79, 160.04, 136.19, 135.46, 128.60, 128.52, 94.41, 67.06, 53.26. HRMS (M + H$^+$): Expected: 291.0756, Found: 291.0744.

20200115_snap-n3_PROTON_001

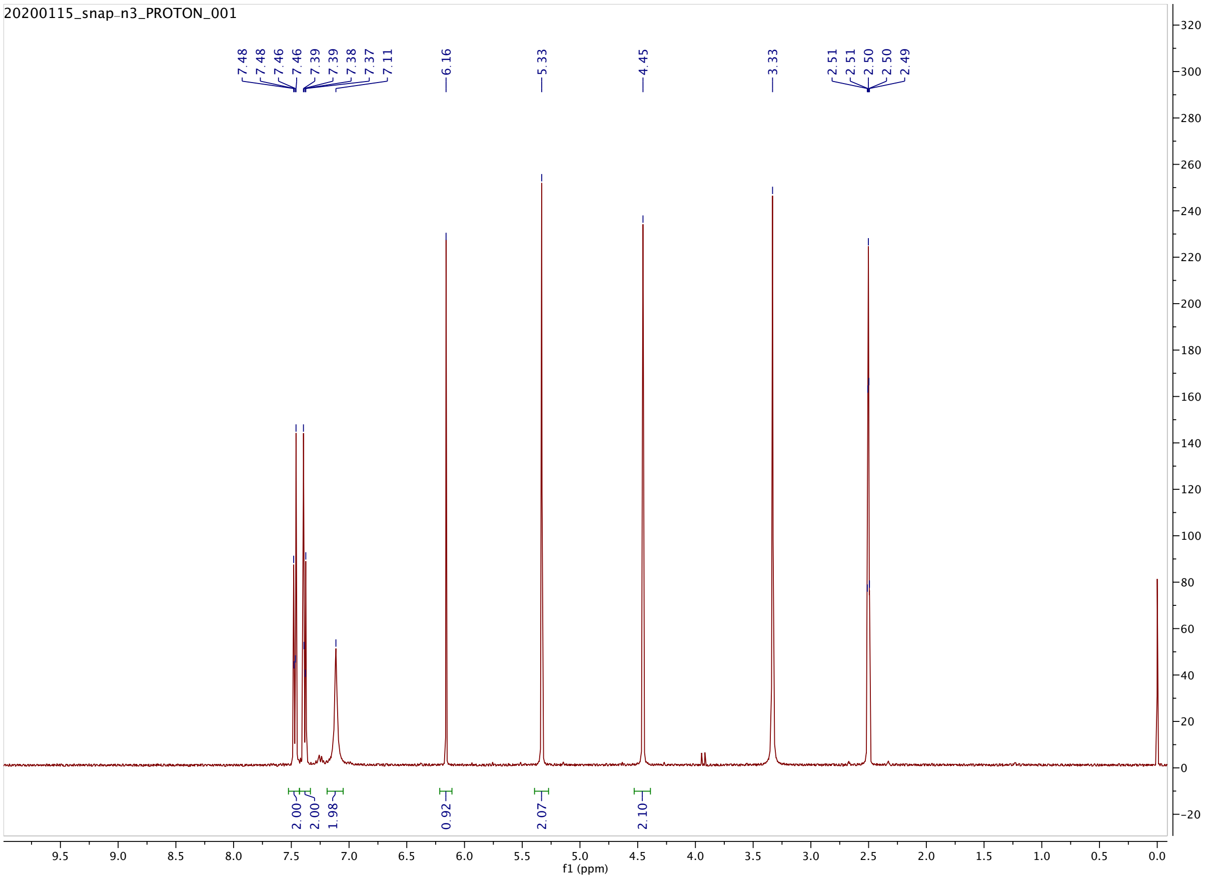

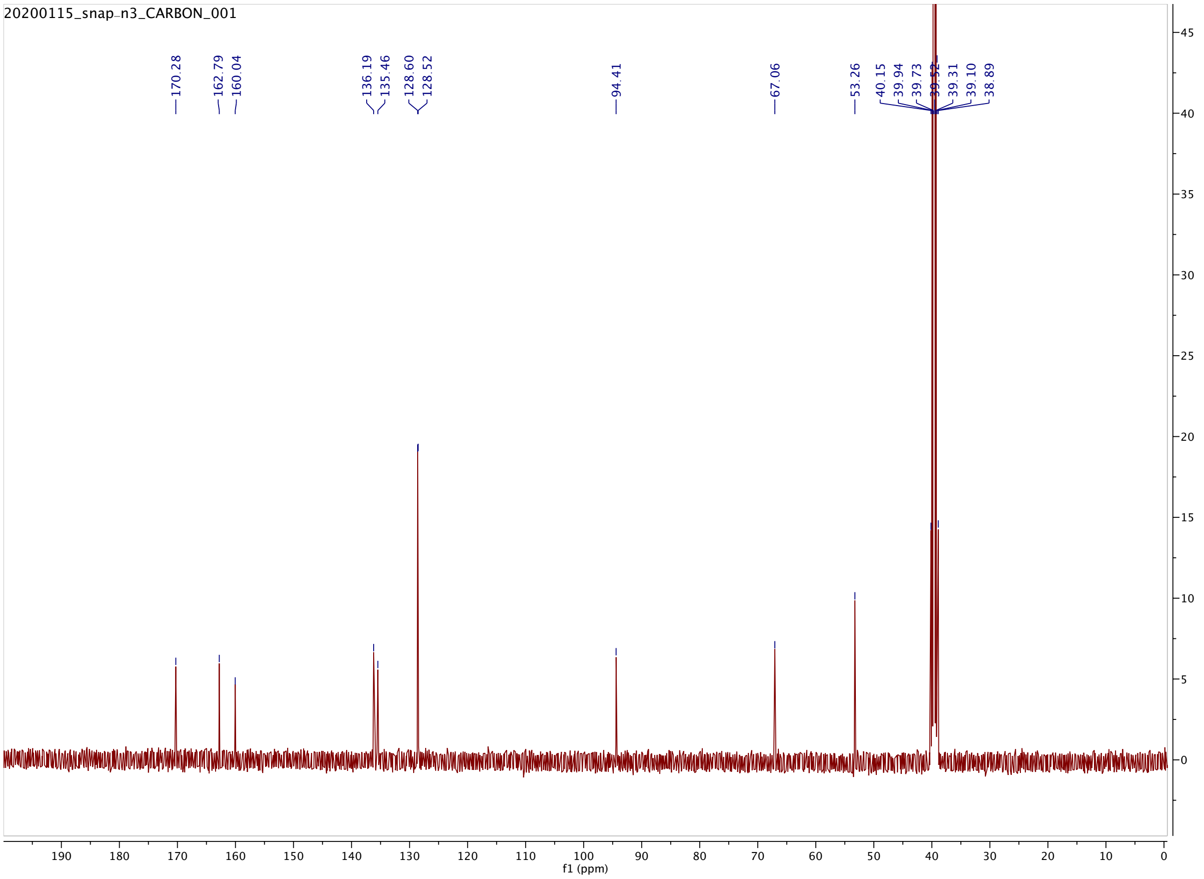
20200115_snap-n3_CARBON_001
170.28
162.79
160.04
136.19
135.46
128.60
128.52
94.41
67.06
53.26
40.15
39.94
39.73
39.52
39.31
39.10
38.89
f1 (ppm)

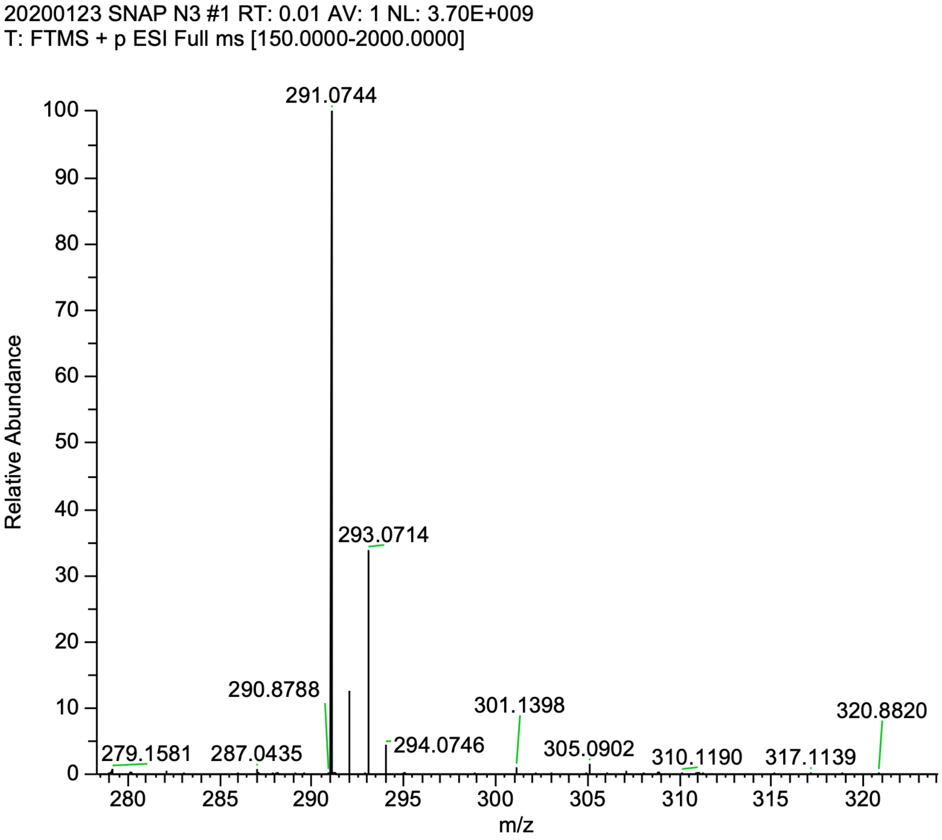


**4-((4-(aminomethyl)benzyl)oxy)-6-chloropyrimidin-2-amine (SNAP-$NH_2$). 2** (500 mg, 1.72 mmol) was dissolved into THF (60 mL) with triphenylphospine (903 mg, 3.45 mmol). Three mL of $H_2O$ were added, and the reaction was allowed to stir overnight at room temperature. The following day, all solvent was removed using a rotary evaporator and the residue was further dried under vacuum. The solid was suspended into cold DCM (~15 mL) and filtered to remove triphenyl phosphine oxide, yielding a fluffy white solid (243 mg, 57% yield).

$^1$H NMR (400 MHz, DMSO-$d_6$) δ 7.34 (q, $J$ = 8.3 Hz, 4H), 7.10 (s, 2H), 6.13 (s, 1H), 5.28 (s, 2H), 3.70 (s, 2H), 1.82 (s, 2H). $^{13}$C NMR (100 MHz, DMSO-$d_6$) δ 170.34, 162.79, 159.99, 144.32, 133.96, 128.26, 127.04, 94.42, 67.38, 45.40. HRMS (M + $H^+$): Expected: 265.0851, Found: 265.0842.

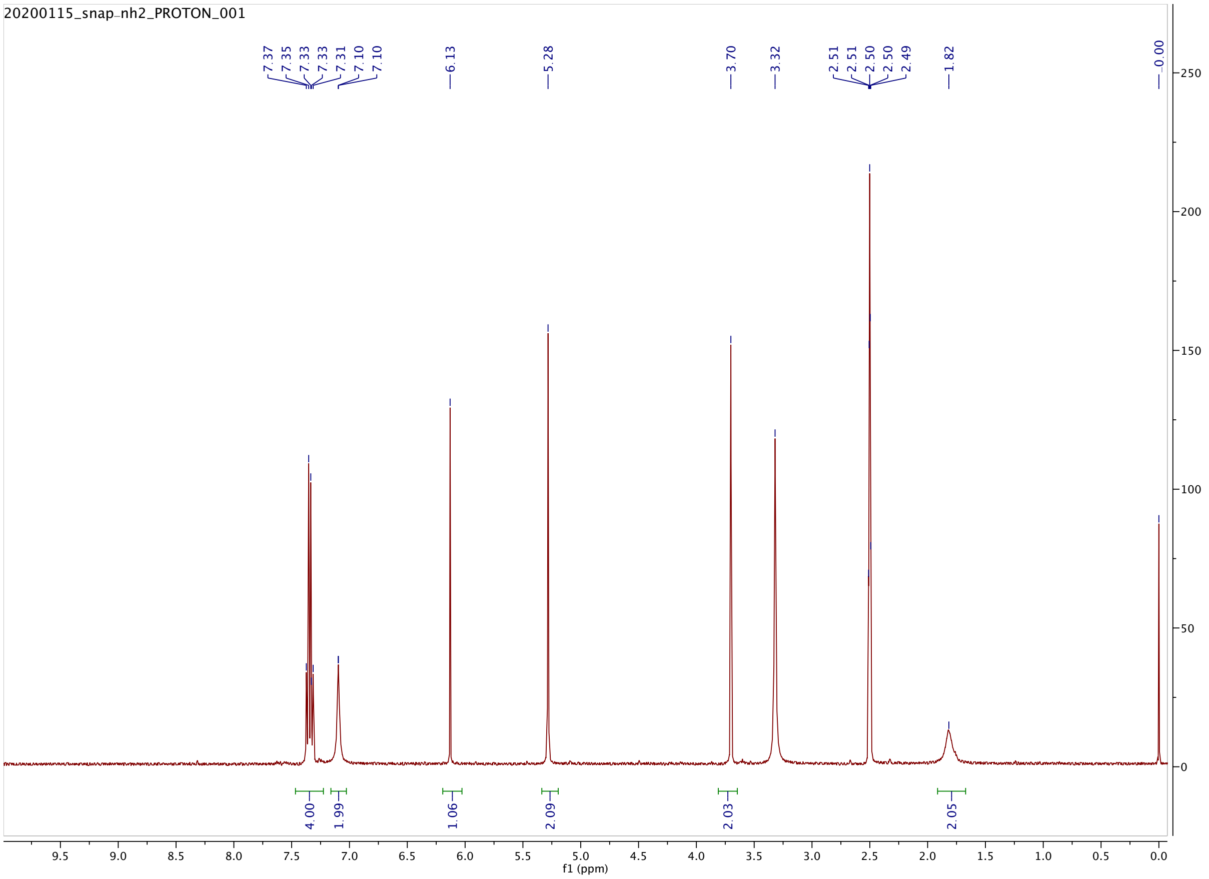

20200115_snap-nh2_PROTON_001
7.37
7.35
7.33
7.33
7.31
7.10
7.10
6.13
5.28
3.70
3.32
2.51
2.51
2.50
2.50
2.49
1.82
0.00
4.00
1.99
1.06
2.09
2.03
2.05
f1 (ppm)

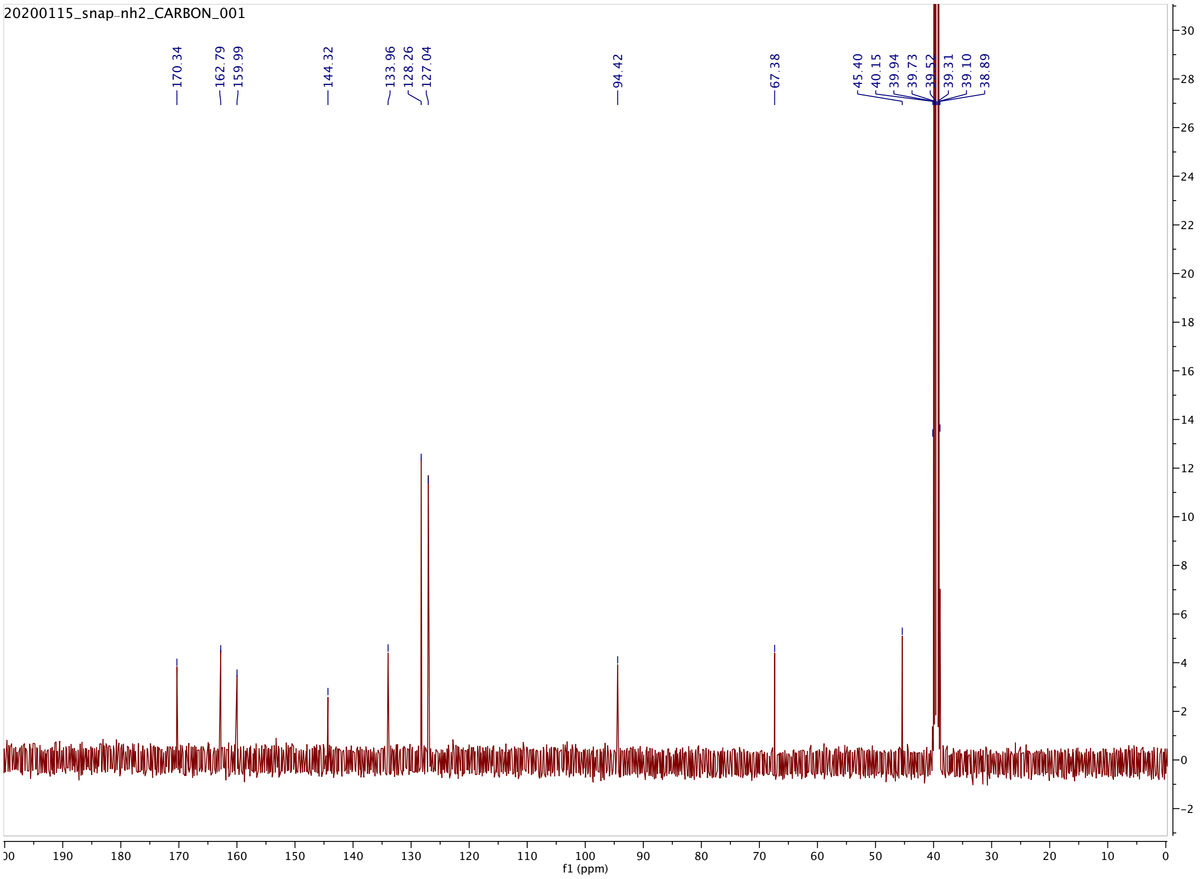
20200115_snap-nh2_CARBON_001
170.34
162.79
159.99
144.32
133.96
128.26
127.04
94.42
67.38
45.40
40.15
39.94
39.73
39.52
39.31
39.10
38.89
f1 (ppm)

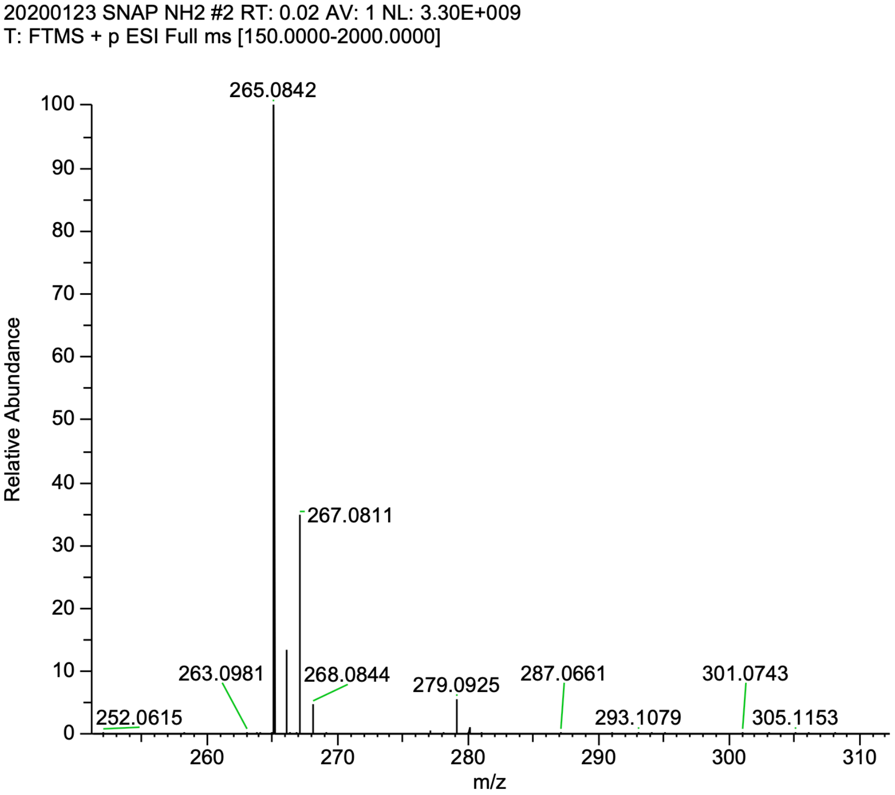
20200123 SNAP NH2 #2 RT: 0.02 AV: 1 NL: 3.30E+009
T: FTMS + p ESI Full ms [150.0000-2000.0000]
265.0842
267.0811
263.0981
268.0844
279.0925
287.0661
301.0743
252.0615
293.1079
305.1153
Relative Abundance
m/z

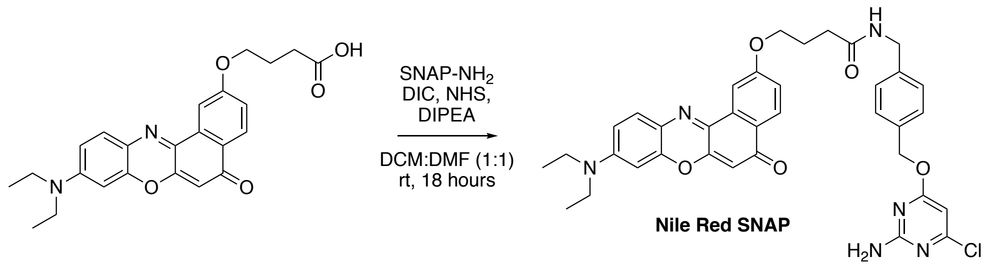

**Nile Red SNAP** was prepared by coupling SNAP-$NH_2$ to Nile Red derivatized with a carboxylic acid handle at the 9-position. Carboxy-Nile Red was prepared using a previously reported synthesis.[1] 3.5 mg (8.3 µmol) of Nile Red-CO2H was coupled to SNAP-$NH_2$ (2.6 mg, 10 µmol) using DIC (1.6 µL, 10 µmol), NHS (1.4 mg, 12.5 µmol), and DIPEA (2.9 µL, 16.7 µmol), and was allowed to stir in a 1:1 mixture of DCM (1 mL) and DMF (1 mL) overnight. The next day, the reaction was diluted into DCM, and washed with 10% citric acid, saturated $NaHCO_3$ and brine. Then the organic layer was dried with $MgSO_4$. The crude product was dried onto silica gel and purified using a gradient of 0-5% MeOH in DCM. A second column using a gradient of 10-100% EtOAc in Hexanes was necessary to completely purify the dye for subsequent photophysical characterization and cellular studies. After purification, 3.2 mg of NR-SNAP was obtained (58% yield).

$^1$H-NMR ($CDCl_3$ + Trace MeOD for solubility, 400 MHz): δ 8.15 (d, *J* = 8.7 Hz, 1H), 7.99 (d, *J* = 2.6 Hz, 1H), 7.58 (d, *J* = 9.1 Hz, 1H), 7.26 – 7.16 (m, 4H), 7.06 (dd, *J* = 8.7, 2.6 Hz, 1H), 6.65 (dd, *J* = 9.1, 2.7 Hz, 1H), 6.44 (d, *J* = 2.7 Hz, 1H), 6.28 (s, 1H), 6.07 (s, 1H), 5.18 (s, 2H), 4.41 (d, *J* = 5.2 Hz, 2H), 4.19 (t, *J* = 5.8 Hz, 2H), 3.46 (q, *J* = 7.1 Hz, 4H), 2.48 (t, *J* = 7.1 Hz, 2H), 2.22 (p, *J* = 6.2 Hz, 2H), 1.25 (t, *J* = 7.0 Hz, 6H). $^{13}$C-NMR ($CDCl_3$ + Trace MeOD for solubility, 100 MHz): δ 183.59, 172.58, 172.50, 170.88, 161.63, 160.85, 152.34, 151.04, 147.01, 139.64, 138.38, 135.32, 134.20, 131.21, 128.49, 128.03, 127.86, 125.71, 124.88, 118.36, 109.89, 106.66, 105.17, 97.23, 96.36, 67.96, 67.26, 45.22, 43.32, 32.90, 25.22, 12.71. HRMS (M + $H^+$): Expected: 667.2430, Found: 667.2429

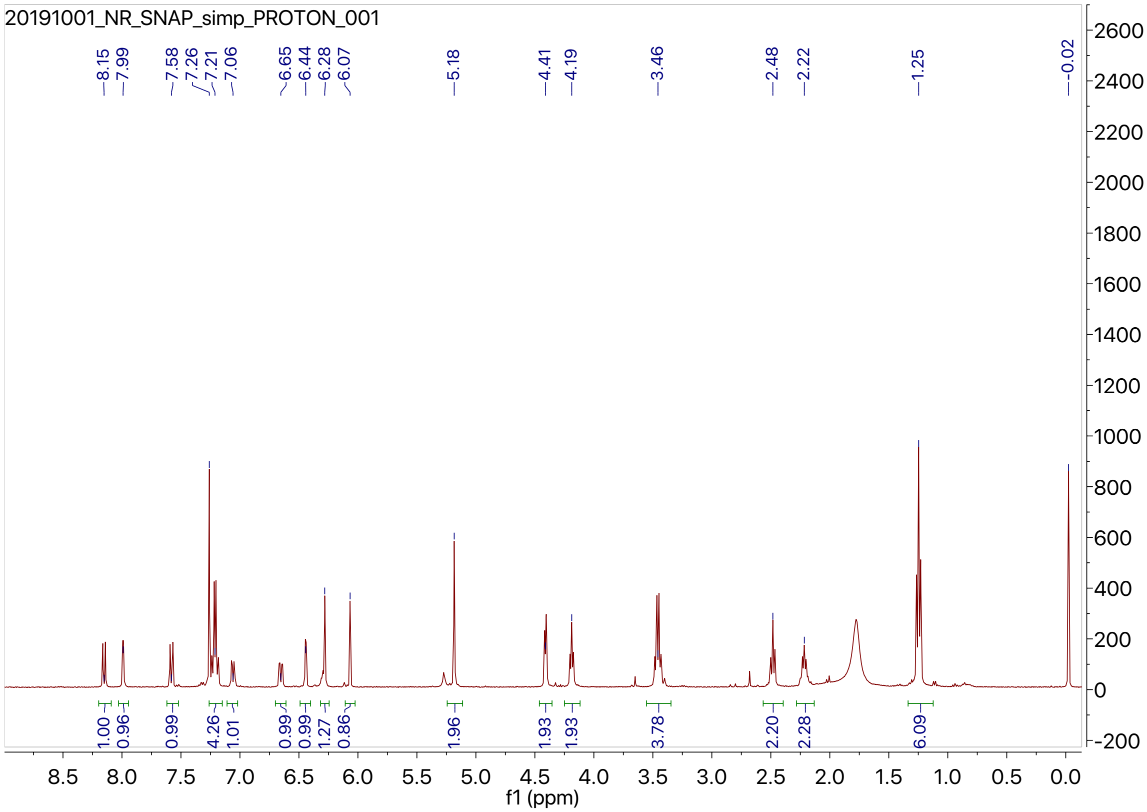
20191001_NR_SNAP_simp_PROTON_001
8.15
7.99
7.58
7.26
7.21
7.06
6.65
6.44
6.28
6.07
5.18
4.41
4.19
3.46
2.48
2.22
1.25
-0.02
1.00
0.96
0.99
4.26
1.01
0.99
0.99
1.27
0.86
1.96
1.93
1.93
3.78
2.20
2.28
6.09
f1 (ppm)

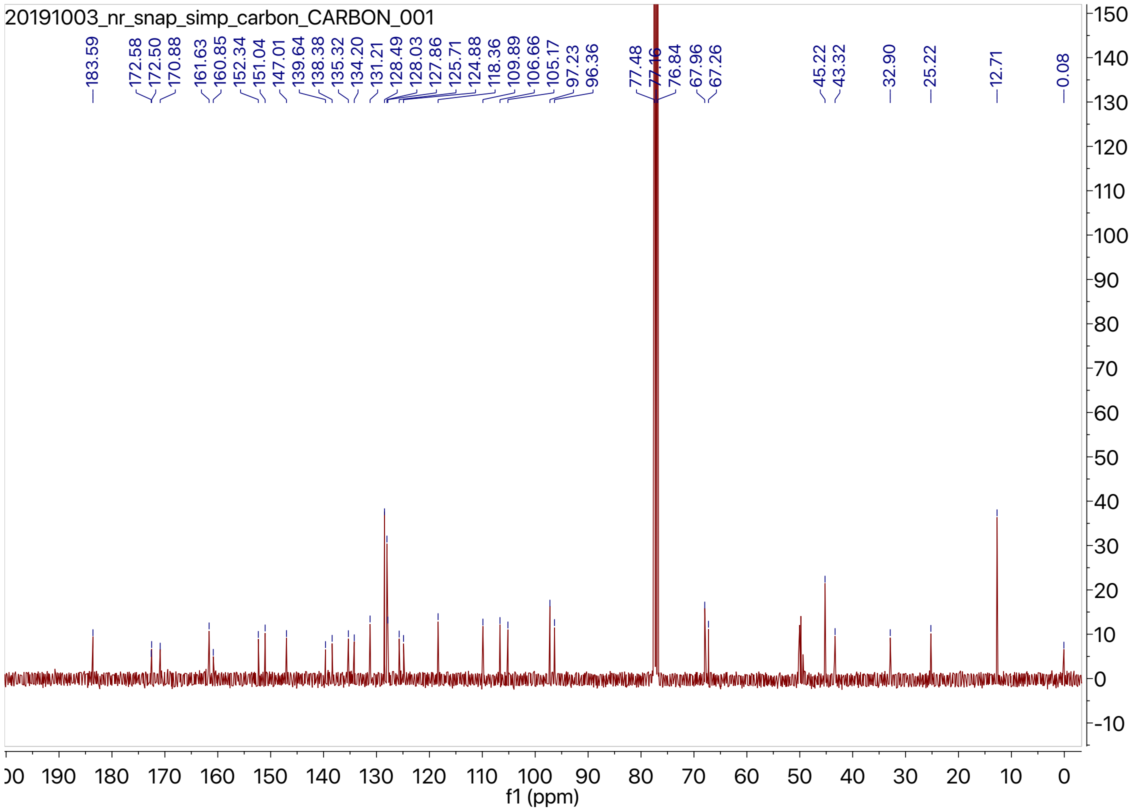
20191003_nr_snap_simp_carbon_CARBON_001
183.59
172.58
172.50
170.88
161.63
160.85
152.34
151.04
147.01
139.64
138.38
135.32
134.20
131.21
128.49
128.03
127.86
125.71
124.88
118.36
109.89
106.66
105.17
97.23
96.36
77.48
77.16
76.84
67.96
67.26
45.22
43.32
32.90
25.22
12.71
0.08
f1 (ppm)

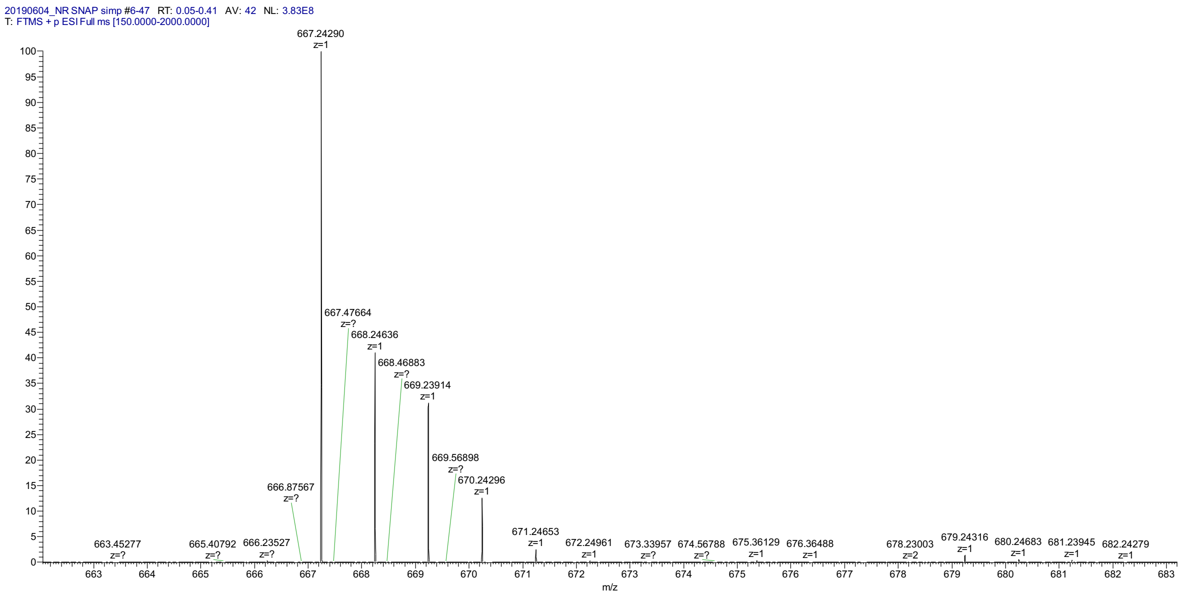


**Supplemental Table 1.** Photophysical properties of Nile Red SNAP.

| Solvent | Abs λmax (nm) | ε (M-1cm-1) | Ex/Em λmax (nm) | Quantum Yield (%) | Brightness (QY x ε) |
|---|---|---|---|---|---|
| **Nile Red SNAP** | | | | | |
| DMSO | 551 | 46,000 ± 3,300 | 549/625 | 78 ± 2 % | 36,000 |
| MeOH | 557 | 50,000 ± 3,600 | 555/632 | 52 ± 1 % | 26,000 |
| BuOH | 548 | 44,000 ± 2,700 | 549/625 | 67 ± 2 % | 29,000 |
| Dioxane | 519 | 43,000 ± 3,700 | 518/581 | 74 ± 2 % | 32,000 |
| H2O | 537* | 29,000 ± 1,400 | 598/655 | n.d. | n.d. |

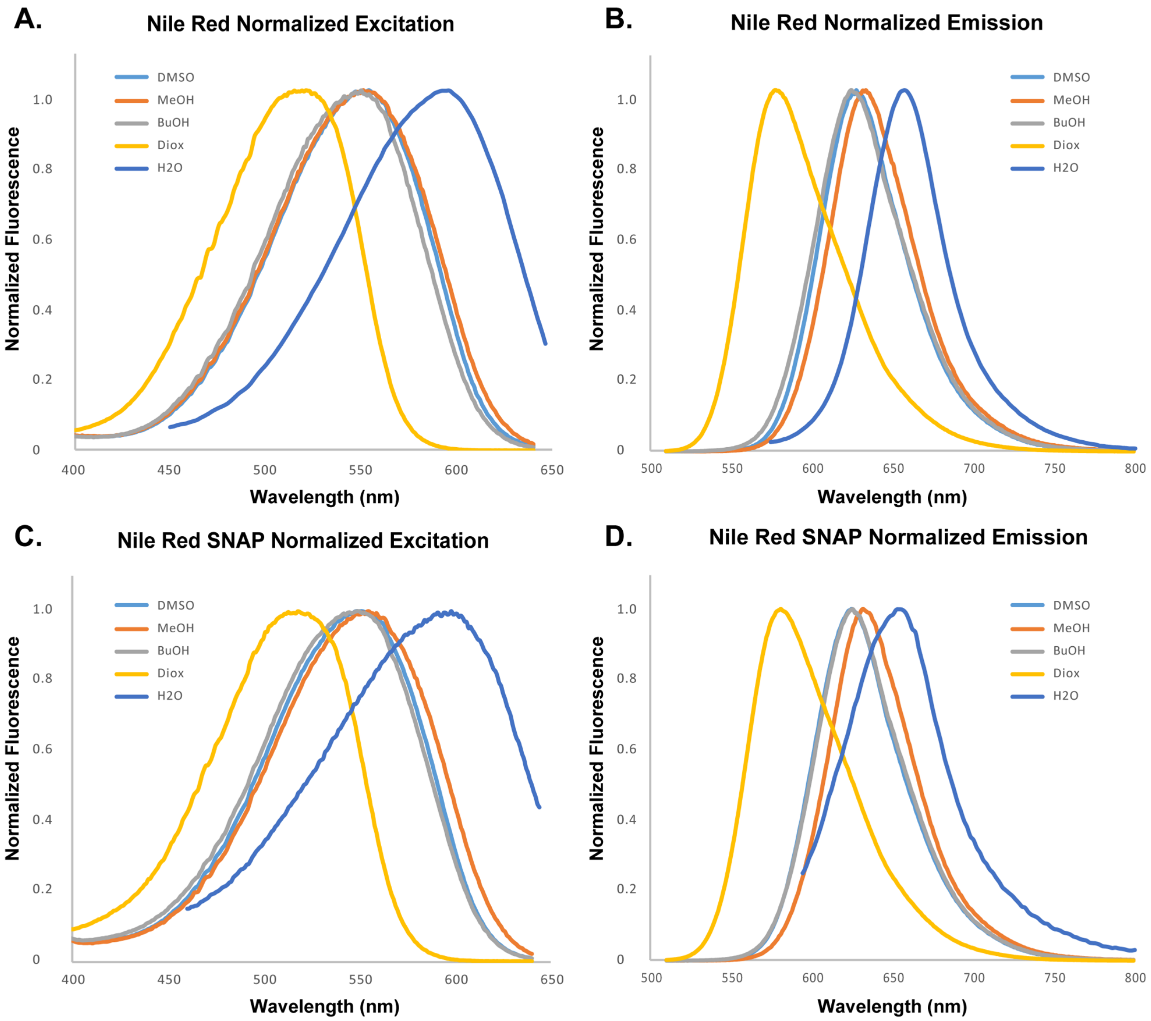


**Supplemental Figure 1.** Normalized Excitation and Emission spectra of Nile Red (A, B) and Nile Red SNAP (C, D) in various solvents.

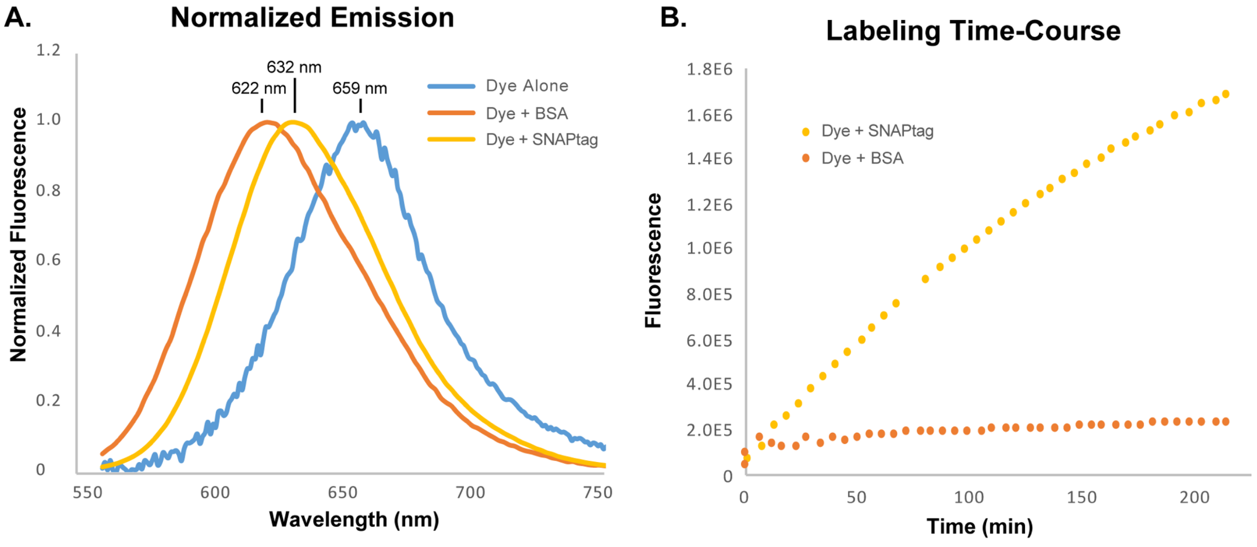

**Supplemental Figure 2.** Response of NR SNAP to addition of BSA or purified SNAP-tag protein. 500 nM dye in PBS containing 1 mM DTT was combined with a 5-fold excess of protein and was allowed to incubate at room temperature. (A) Normalized spectra show the emission changes produced by labelling SNAP-tag versus non-specific interaction with BSA. (B) Fluorescence showed a time-dependent increase during SNAP-tag labelling, while the non-specific interaction with BSA only induced a small signal increase after mixing.

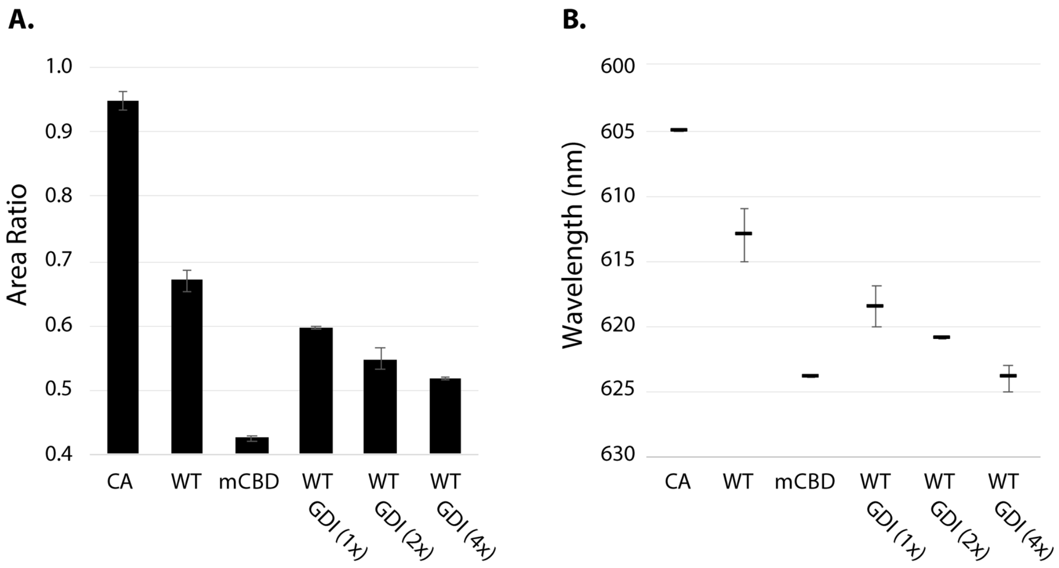


**Supplemental Figure 3.** Optimization of RhoGDI expression level. Increasing amounts of GDI decrease the response of the sensor towards the mCBD negative control, but cell health decreases as more GDI is added. Though the sensor response could likely be decreased to that of the unbound negative control, we could not express higher than a 4x excess without compromising cell health to the degree that the expression level of the sensor became difficult to measure.

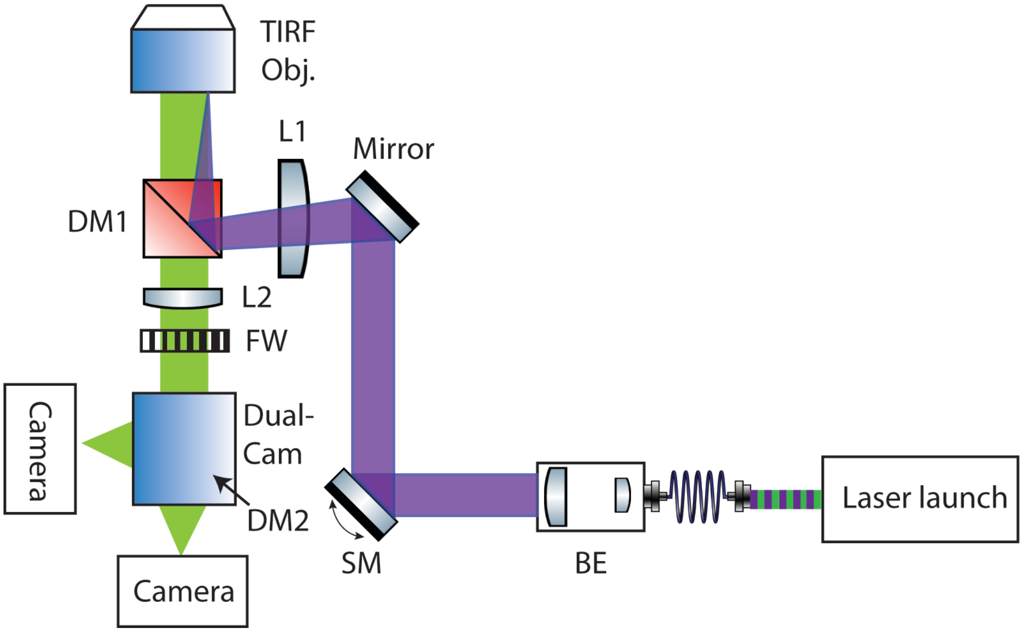


**Supplemental Figure 4.** Scope setup for ratiometric imaging of SNAPsense biosensor using NR SNAP.

**Cloning.** The original SNAP biosensor construct was created by modifying a single-chain biosensor for Cdc42 designed to be expressed using UAA mutagenesis. The plasmid for this sensor is in a pTriEx backbone and contains CBD, sfGFP, and Cdc42 separated by flexible spacers. In this plasmid, CBD contained a mutation at F271 to the amber stop codon (TAG) for incorporation of an unnatural amino acid. We first mutated this site back to the native phenylalanine, then replaced sfGFP with SNAP-tag through restriction cloning using 5' EcoRI and 3' NotI restriction sites. For all GSG spacers, the DNA encoding these spacers was incorporated into the forward and reverse primers when amplifying SNAP for restriction insertion. All other mutations were carried out using Q5 site-directed mutagenesis (NEB). During optimization, the CBD-SNAP-Cdc42CA and mCBD-SNAP-Cdc42CA variants of each mutant were tested.

To enable Nile Red brightness changes associated with Cdc42 activation to be measured, sfGFP was incorporated at the N-terminus of the plasmid using Gibson assembly. Stable cell lines were created by cloning the optimized construct into the piggybac backbone.[2] Again, restriction cloning was used to insert the DNA encoding the sensor into a piggybac plasmid using 5' AgeI and 3' SalI restriction sites. Biosensor expression was controlled by doxycycline using a TetOFF system, and plasmids were created for stable cell line generation using either Hygromycin or Puromycin.

All plasmids were verified by sequencing (Genewiz, South Plainfield, NJ) before use. Primer sequences used for all cloning steps are available upon request.

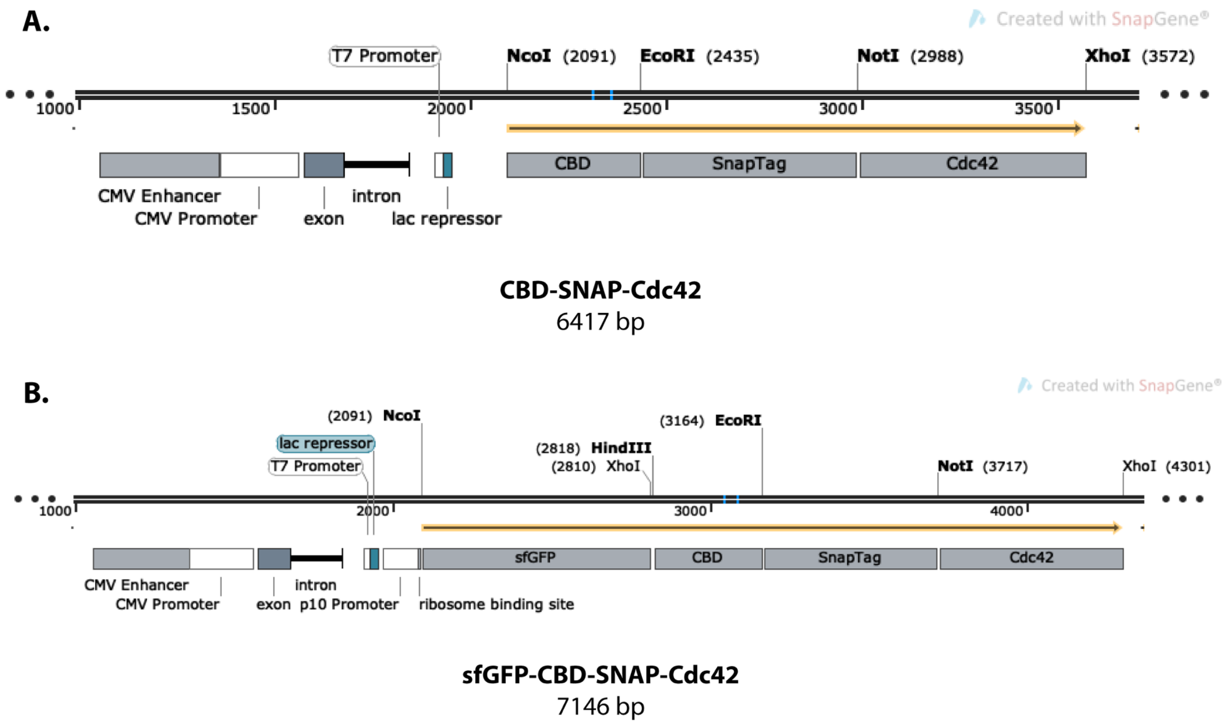


**Supplemental Figure 5.** General plasmid maps for the coding regions of the SNAPsense Cdc42 biosensors (pTriEx backbone). For simplicity, only restriction sites used for cloning are shown.

**Cell culture**. Mouse embryonic fibroblasts (MEF, Clontech) and HeLa cells were maintained in 10% $CO_2$ at 37 °C in culture medium composed of Dulbecco's modified Eagle's medium (DMEM, Cellgro) with 10% fetal bovine serum (HyClone, Thermo Scientific) and 2 mM GlutaMax (Gibco, Life Technologies).

The biosensor constructs were inserted into a tet-off inducible piggybac system and stable lines were produced in tet-off MEF cells.[3] The stable MEF cells were cultured in medium supplemented with 1 µg/mL doxycycline to suppress biosensor expression. Doxycycline was removed three days before imaging. Before imaging, the cells were incubated with 100 nM Nile red dye for 1 hour and washed 3 times with the imaging buffer (Ham's F-12, Kaighn's Modification (Caisson Laboratories, Inc), 15 mM HEPES and 5% FBS).

**Transfection, Dye Labeling, and Fluorescence Measurements.** HeLa cells were plated into 6- or 12-well plates at a density of 40,000 cells or 20,000 cells per well, respectively. The following day, media was exchanged with fresh media containing 100 nM Nile Red SNAP (2 mL/well for 6-well dish, 1 mL/well for 12-well disn), and the cells were transfected with plasmid DNA using Fugene 6 (0.5 µg DNA/well of a 6-well dish, 0.25 µg DNA/well of a 12-well dish, 3:1 µL reagent/µg DNA). After another overnight incubation, the media was exchanged with dye-free media (1 or 2 mL/well) and cells were allowed to incubate for 1 hour to remove excess dye. The cells were then washed with PBS, trypsinized (200 uL/well of a 6-well plate, 150 uL/well of a 12-well plate), diluted with 4x medium, and each well was transferred into a separate Eppendorf tube. Cells were pelleted by centrifuging at 600 rcf for 5 minutes, the media was carefully aspirated off, and the cells were resuspended into PBS (500 mL if from 6-well plate, 400 mL if from 12-well plate). At this point, the emission spectrum of Nile Red was directly measured in the suspension cells.

**Biosensor imaging**. Biosensor imaging was performed on a home-built total internal reflection fluorescence microscope, equipped with four solid-state lasers (Coherent OBIS 405 nm, 488 nm, 561 nm and 647 nm). A four-band dichroic mirror (DM: Di01-R405/488/561/635, Semrock) was used for multi-color imaging. Fluorescence images were collected using a 60 X TIRF objective (UAPON 60XOTIRF, NA 1.45, Olympus) and split by a dichroic mirror (Semrock, FF624-Di01) through a dual-camera system (Hamamatsu, W-VIEW GEMINI-2C). The images were projected onto two metal-oxide-semiconductor (sCMOS) cameras (Photometric, 95B Prime). The laser power was set to 1 mW and the exposure time was set to 100 ms with 3 seconds interval between time points.

JF646 MEFs were imaged using an inverted IX81 epifluorescence microscope (Olympus) with a 40x 1.3NA Silicon oil objective (UPLSAPO40XS, Olympus) and a ORCA-Flash4.0 V2+ sCMOS camera (Hamamatsu). Images were collected every 10 sec for a total of 15 min using FF405/496/593/649-Di01 as a dichroic (Semrock), HQ630/40X and HQ470/40X (Chroma) as excitation filters and ET705/72 and HQ525/50M (Chroma) as emission filters (for JF646 and GFP respectively).

**Biosensor image processing**. Biosensor image processing was as previously described.[2] Briefly, the raw images were passing through a processing pipeline including shade correction, baseline subtraction, masking, registration, and photo-bleach correction when needed, then ratiometric images were obtained by dividing two images. The range of the color bar was set to the 5th and 95th percentiles of the ratio images. The correlation analysis between Cdc42 activity and edge velocity was performed following using a previously published protocol.[4]
